# A SMART method for efficiently isolating monoclonal antibodies from individual rhesus macaque memory B cells

**DOI:** 10.1101/2023.06.02.543510

**Authors:** Jason T. Weinfurter, Sarah N. Bennett, Matthew Reynolds

## Abstract

Characterizing antigen-specific B cells is a critical component of vaccine and infectious disease studies in rhesus macaques (RMs). However, it is challenging to capture immunoglobulin variable (IgV) genes from individual RM B cells using 5’ multiplex (MTPX) primers in nested PCR reactions. In particular, the diversity within RM IgV gene leader sequences necessitates the use of large 5’ MTPX primer sets to amplify IgV genes, decreasing PCR efficiency. To address this problem, we developed a switching mechanism at the 5’ ends of the RNA transcript (SMART)-based method for amplifying IgV genes from single RM B cells, providing unbiased capture of Ig heavy and light chain pairs for cloning antibodies. We demonstrate this technique by isolating simian immunodeficiency virus (SIV) envelope-specific antibodies from single-sorted RM memory B cells. This approach has several advantages over existing methods for PCR cloning antibodies from RMs. First, optimized PCR conditions and SMART 5’ and 3’ rapid amplification of cDNA ends (RACE) reactions generate full-length cDNAs from individual B cells. Second, it appends synthetic primer binding sites to the 5’ and 3’ ends of cDNA during synthesis, allowing for PCR amplification of low-abundance antibody templates. Third, universal 5’ primers are employed to amplify the IgV genes from cDNA, simplifying the primer mixes in the nested PCR reactions and improving the recovery of matched heavy and light chain pairs. We anticipate this method will enhance the isolation of antibodies from individual RM B cells, supporting the genetic and functional characterization of antigen-specific B cells.

## Introduction

Rhesus macaques (RMs) are valuable large-animal models for studying human pathogens, providing insights into protective immunity and pre-clinical tests of vaccine efficacy. Since antibodies are the primary mediator of humoral immune responses, monoclonal antibodies (mAbs) are often isolated to investigate their specificity, function, and variable (V) gene usage. However, cloning antibodies from RMs is challenging. Although phage-display and B-cell immortalization platforms have been used to isolate mAbs from RMs (Holman et al., 2017; Kuwata, Katsumata, Takaki, Miura, & Igarashi, 2011; Samsel et al., 2023; Blasi et al., 2018), these approaches are laborious and often yield relatively few antigen-specific antibodies. Alternatively, PCR-based strategies aim to amplify heavy and light chain genes directly from individual B cells, then produce recombinant antibodies in mammalian cell lines (Magnani et al., 2017; Meng et al., 2015; Silveira et al., 2015; Sundling et al., 2014; Wiehe et al., 2014).

However, the complexity of RM Ig genes hinder the cloning efficiency of these approaches. PCR-based methods for cloning RM antibodies are built on well-established protocols for isolating mAbs from single-cell sorted human and murine B cells (Tiller et al., 2008; Wrammert et al., 2008; Guthmiller, Dugan, Neu, Lan, & Wilson, 2019; von Boehmer et al., 2016). These cloning strategies use nested PCRs to amplify immunoglobulin variable (IgV) genes. In these schemes, the 5’ primers typically anneal to the Ig leader sequences, which are immediately upstream of the IgV genes and are relatively conserved within human and murine V gene families. Thus, for these species, only small panels of 5’ primers are needed to capture most IgV genes. In contrast, RM heavy and light chain genes have more copy-number variations and diversity than their human or murine orthologs (Ramesh et al., 2017; Vázquez Bernat et al., 2021; Cirelli et al., 2019). To compensate for this increased genetic diversity, large cocktails of multiplex (MTPX) primers are used to amplify RM IgV genes (Sundling, Phad, Douagi, Navis, & Karlsson Hedestam, 2012b; Silveira et al., 2015; Mason et al., 2016; Wiehe et al., 2014; Feng et al., 2023). However, increasing the number of primers can reduce PCR efficiency through competition for template binding and inter-primer interference. Moreover, although recent studies have advanced our understanding of the RM Ig loci (Cirelli et al., 2019; Vázquez Bernat et al., 2021; Chernyshev, Kaduk, Corcoran, & Karlsson Hedestam, 2021; Ramesh et al., 2017), gaps in the RM Ig gene repertoire likely remain. Thus, existing 5’ MTPX primer panels may be incomplete and unable to capture all of the RM IgV genes.

Antigen-specific memory B cells are often used as source material for cloning antibodies. These antigen-experienced cells express ample B-cell receptors (BCRs), allowing the use of fluorescently-labeled antigenic probes to select individual antigen-binding B cells via fluorescent-activated cell sorting (FACS). However, since memory B cells are not actively secreting antibodies, they have a low abundance of antibody transcripts (Phad et al., 2022; Meng et al., 2015; Tiller et al., 2008; Upadhyay et al., 2018), limiting the recovery of heavy and light chain pairs (Meng et al., 2015).

Using PCR-based techniques, the first step in cloning antibodies from individual memory B cells is converting mRNA into cDNA using reverse transcriptase (RT). SMART (switching mechanism at the 5’ ends of the RNA transcript) technology, a refinement of 5’ and 3’ rapid amplification of cDNA ends (5’/3’ RACE) synthesis, is a useful technique for generating full-length cDNA libraries when mRNA transcripts are low (Zhu, Machleder, Chenchik, Li, & Siebert, 2001). SMART technology takes advantage of the natural poly(A) tail of mRNAs and the template-switching activity of Moloney murine leukemia virus (MMLV)-based RTs. During 3’ RACE reactions, cDNA synthesis is initiated using primers containing deoxythymidine (dT) tracts that are complementary to mRNA poly(A) tails. As RTs with transferase activity reach the 5’ cap of mRNAs, they add non-templated nucleotides, predominantly deoxycytidines (+CCC), to the 3’ end of nascent cDNA molecules (Kulpa, Topping, & Telesnitsky, 1997). The newly added +CCC bases can then bind to template-switching oligos (TSOs) with complementary riboguanosine nucleotides (rGrGrG) at their 3’ end (Zhu et al., 2001). This prompts the RT to complete cDNA synthesis using the TSO as the template, effectively switching templates in the process. 5’ RACE is particularly useful when the 5’ ends of transcripts are highly variable and designing primers for downstream amplifications is challenging. In such cases, synthetic primer binding sites can be incorporated into the TSOs and added to the 5’ ends of cDNAs during synthesis. As a result, 5’ universal primers complementary to the primer binding sites can be paired with gene-specific 3’ primers in PCRs to amplify target genes (Zhu et al., 2001).

In this study, we present a novel method for isolating mAbs from individual RM memory B cells. One of the key features of this protocol is the incorporation of synthetic primer binding sites to the 5’ and 3’ ends of cDNA during synthesis, which offers several advantages for rescuing antibodies. First, low-abundance templates can be PCR amplified using 5’ and 3’ universal primers, which yields sufficient material for downstream PCRs. Second, appending two synthetic primer binding sites into the 5’ end of the cDNAs enables using successive 5’ universal primers in nested PCR reactions, thus eliminating the need for complex 5’ MTPX primer sets to amplify the IgV genes. We demonstrate this process by isolating mAbs specific for the envelope (Env) protein of simian immunodeficiency virus (SIV) from single-cell sorted rhesus macaque memory B cells. We anticipate this approach will improve the recovery of monoclonal antibodies from RMs and facilitate characterizing antibody specificity and functions in pre-clinical studies.

## Materials and Methods

### 2.1 Isolation of SIV Env-specific memory B cells

We isolated SIV-Env-binding memory B cells as previously described (Mohanram et al., 2014). Briefly, we used cryopreserved peripheral blood mononuclear cells (PBMCs) from an Indian-origin rhesus macaque infected with SIVmac239 (Mudd et al., 2012). The banked cells were placed in a 37°C water bath until just thawed, washed twice with 10ml of prewarmed R10 (RPMI 1640 media supplemented with 10% fetal bovine serum), and resuspended in 100 µl of R10. Afterward, the cell suspensions were incubated with antibodies against CD4 (unlabeled; OKT4 and 19Thy5D7), CD3 (FITC; SP34), CD14 (FITC; M5E2), CD16 (FITC; 3G8), CD20 (PerCP-Cy5.5; 2H7), IgM (PE-Cy7; MHM-88), CD27 (APC; O323), and near-infrared LIVE/DEAD dye for 30 minutes at 4°C. The anti-CD4 antibodies were added to block the non-specific binding of SIV Env. After washing the cells twice with R10, we added 4µg of SIVmac239 Env gp130 (NIH HIV Reagent Program, ARP-12797), which we biotinylated in-house using the Biotin-XX Microscale Protein Labeling Kit (Invitrogen) according to the manufacturer’s instructions, and incubated for 20 minutes at 4°C. The cells were then washed twice in R10, suspended in 50µl of FACS buffer (1XPBS supplemented with 2% FCS), mixed with 50µl of streptavidin-PE diluted 1:400 in FACS buffer, and incubated for 20 minutes at 4°C. Lastly, the cells were washed twice with R10 and suspended at a concentration of 3e6/ml in R10.

Before sorting, the cell suspension was passed through a .45µm filter to remove cell aggregates. Next, single live CD3-, CD14-, CD16-, CD20+, CD27+, IgM-, SIV Env gp130+ lymphocytes (Fig. 1) were sorted into individual wells of a 96-well plate containing 4 µl of lysis buffer using a FACSJazz Cell Sorter running FACSDiva software (Becton Dickinson), leaving one row of the 96-well plate empty as contamination controls for downstream PCRs. The lysis buffer consisted of 0.9 µl of nuclease-free water, 1.9 µl of 0.1% (vol/vol) Triton-X, 1 µl of a 10 mM dNTP mix, 0.1 µl of 100 µM poly(dt) RT oligo, and 0.1 µl of 40U/µl of RNAse inhibitor. The poly(dT) RT oligo (Table 1) has a synthetic primer binding sequence incorporated into the 5’ end, termed 3’ universal (3U), and “VN” nucleotides at the 3’ end, anchoring the oligonucleotide to the beginning of the mRNA poly-A tail as previously described (Picelli et al., 2014; Liao & Gong, 1997). After sorting, the 96-well plate was immediately placed on dry ice and stored at -80°C.

**Table 1.**
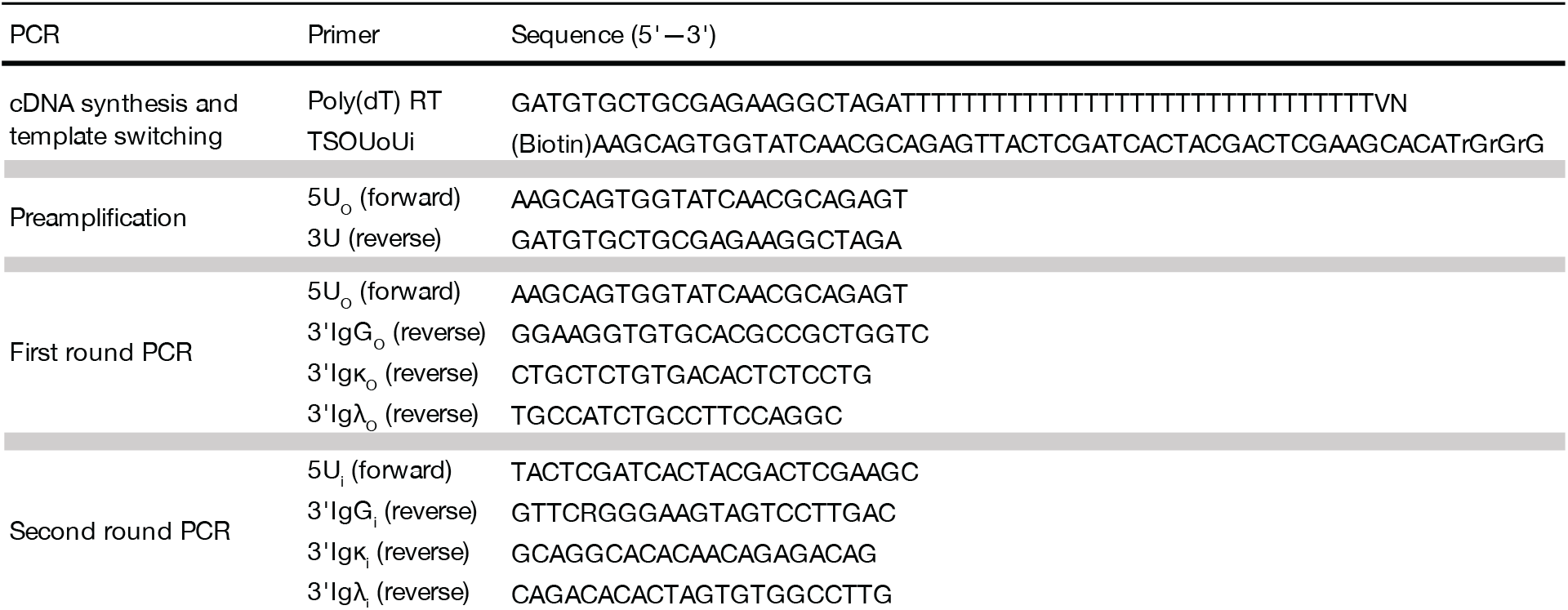
Primers

**Figure 1.**
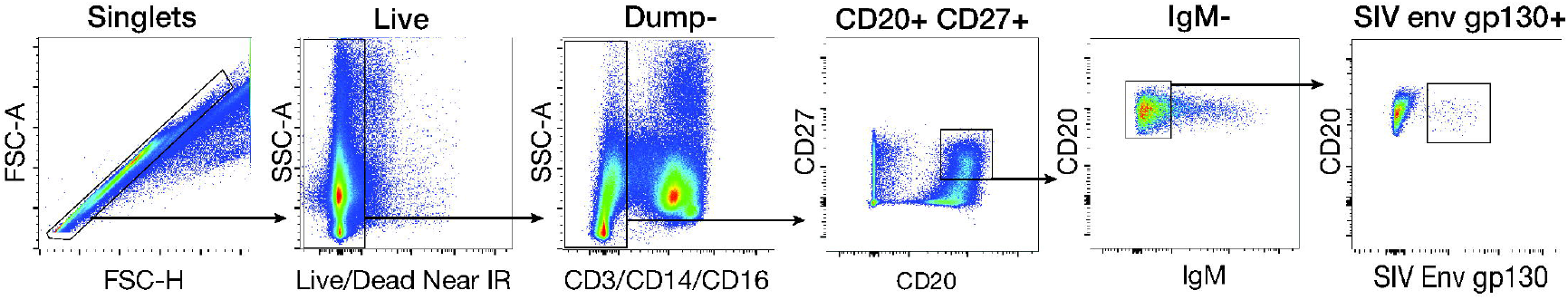
Gating strategy. Representative gating strategy for isolating SIV Env gp130-binding memory B cells (single live CD3-, CD14-, CD16-, CD20+, CD27+, IgM-, SIV Env gp130+ cells) from a RM chronically infected with SIVmac239.

### 2.2 cDNA synthesis and preamplification

To generate cDNA, the 96-well plate containing the sorted cell lysates was thawed on ice and incubated for 3 minutes at 72°C, annealing the poly(dT) RT primer to the mRNA (Fig. 2). To each well, we added 7 µl of the RT mix that contained 0.5 µl of 200 U/µl Superscript IV (Invitrogen), 0.25 µl of 40 U/µl RNAseOUT (ThermoFisher Scientific), 2.2 µl Superscript IV 5X buffer, 0.55 µl of 100 mM dithiothreitol, 2.2 µl of 5 M betaine, 0.11 µl of 100 µM template switching oligo (TSOUoUi), and 1.19 µl of nuclease-free water. Of note, the TSOUoUi oligo has two synthetic primer binding sites incorporated, termed 5’ universal outer (5UO) and 5’ universal inner (5Ui) (Table 1). After briefly centrifuging the samples, we performed the first-strand cDNA synthesis reaction in a thermocycler under the following conditions: 30 minutes at 50°C, 5 minutes at 85°C, and cooling to 4°C. Next, to increase the cDNA yield, we performed a preamplification step using a PCR mixture consisting of 11µl of the first-strand cDNA synthesis product, 12.5 µl of KAPA HiFi HotStart Ready Mix (Roche), 0.03 µl of 100 µM 5UO and 3U primers, and 1.44 µl of nuclease-free water for a total volume of 25µl. The PCR program was as follows: 1 minute at 98°C, 20 cycles of 98°C for 20 seconds, 67°C for 15 seconds, 72°C for 6 minutes, followed by a final elongation step at 72°C for 5 minutes before cooling to 4°C.

**Figure 2.**
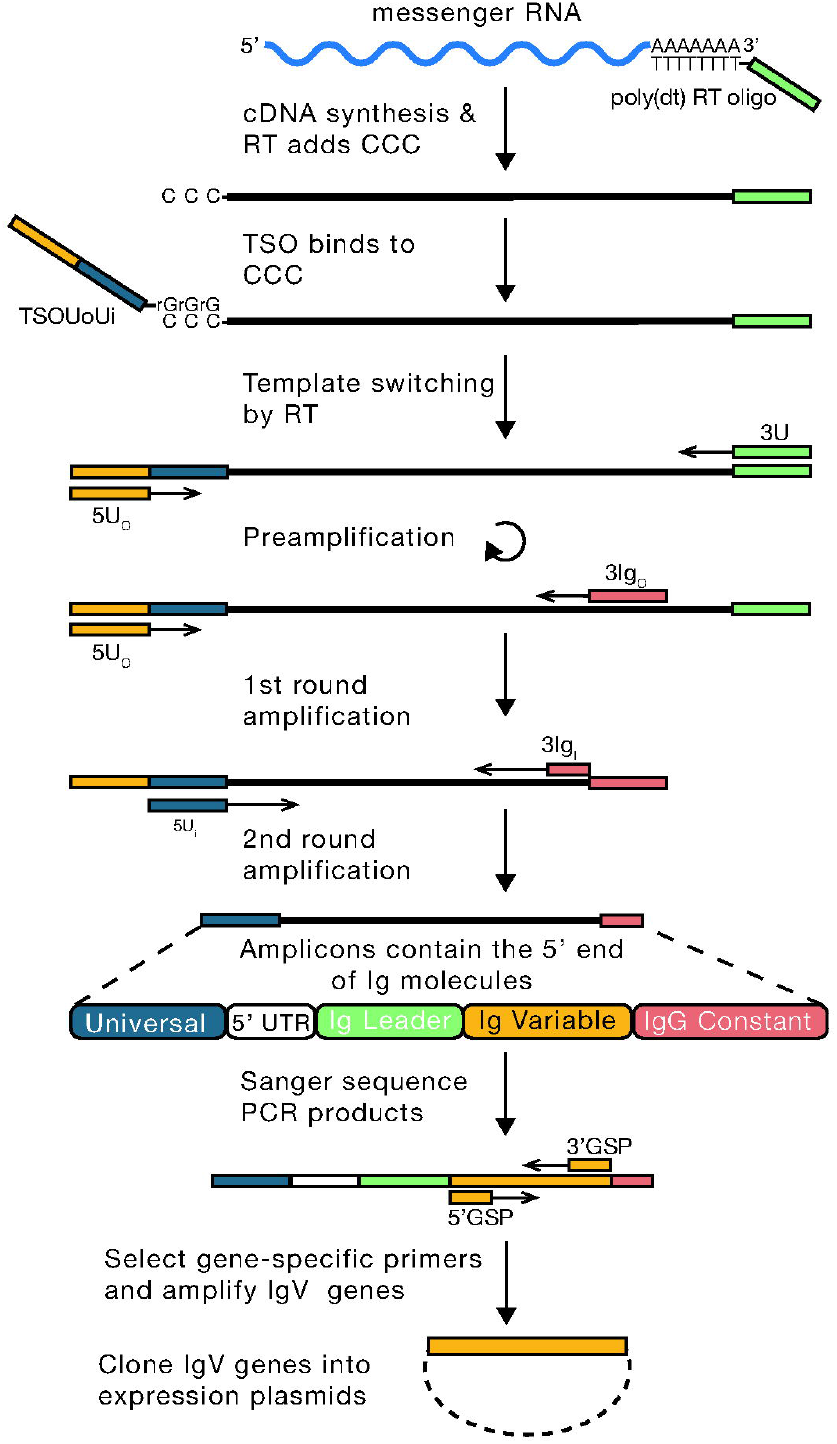
Workflow for cloning antibodies from single cells using cDNA synthesized by 5’RACE. Full-length cDNA is synthesized from the mRNA of individual B cells using an MMLV-based reverse transcriptase (RT) and primers containing synthetic primer binding sites. cDNA synthesis is initiated by the poly(dt) RT oligo, and the RT adds non-templated deoxycytidines (CCC) to the 3’ end of the cDNAs. The template-switching oligo (TSOUolli) then binds to the CCC overhang, and the RT completes cDNA synthesis using TSOUoUi as the template. The resulting cDNAs contain two universal primer binding sites at the 5’ end (5U_o_ and 5U_i_) and one at the 3’ end (3U). A preamplification step with the 5U_o_ and 3U primers is included to increase the amount of templates available for subsequent steps. The first round of PCRs are performed with the universal outer forward primer (5U_o_) and a reverse primer (3Ig_o_) specific for the IgH, IgK, or IgL chain. The second round PCR is performed with the universal inner forward primer (5U_i_) and a nested reverse primer (31g_i_) specific for the IgH, IgK, or IgL chain. The result is a PCR product containing the 5’ end of the Ig templates. These products are sequenced to select gene-specific primers that contain restriction sites to amplify the IgV genes and clone them into expression vectors.

### 2.3 Bead purification of the preamplification PCR products

We purified the preamplification PCR products before amplifying the heavy and light chain variable regions. First, the PCR plate was briefly centrifuged and 25 µl of room-temperature AMPure XP beads (1:1 ratio; Beckman Coulter) was added to each well and incubated for 8 minutes at room temperature. Next, the AMPure XP beads were concentrated by placing the plate on a magnetic rack for 5 minutes. Afterward, ∼45 µl of the supernatant was removed and disposed of. While still on the magnet, each well was washed twice with 200 µl of 80% ethanol, incubating for 30 seconds before removing and disposing of the ethanol. After the last wash, the plate was air-dried for 5 minutes. Next, we removed the plate from the magnetic rack, added 23 µl of nuclease-free water to each well, resuspended the beads with a pipette, and incubated for 2 minutes. The plate was then placed back onto the magnetic rack and incubated for 2 minutes before transferring 20 µl of the supernatant containing the preamplification product to a new 96-well plate, avoiding moving the beads.

### 2.4 Amplification of Ig variable sequences

We focused on amplifying Ig genes from 16 wells (14 with cells and two PCR controls) from the plate prepared above. In nested PCR reactions, we amplified the heavy and light chain (kappa/lambda) variable genes using 5UO and 5Ui oligos complementary to synthetic primer binding sites at the 5’ ends of the cDNAs and 3’ reverse oligos complementary to Ig constant domains (Fig. 2, Table 1). For each reaction, the PCR mixes consisted of 2.5 µl of 10X PCR buffer, 0.5 µl of 10mM dNTP mix, 0.5 µl of 10 µM forward and reverse primers, 1 µl of cDNA, 1.25 U of Hotstar Taq DNA polymerase (Qiagen), and 19.75 µl of water for a total volume of 25 µl. Touchdown PCR conditions were used to increase the specificity of the reactions by reducing the annealing temperature in each of the first ten cycles by 1°C, from 65°C to 55°C. Thus, the PCR programs were initiated with a denaturing step at 95°C for 15 minutes followed by 10 cycles of 30 seconds at 94°C, 30 seconds at 65°C (−1°C per cycle), 1 minute at 72° C, and then 25 cycles of 30 seconds at 94°C, 30 seconds at 55°C, 1 minute at 72°C, and a final elongation step for 10 minutes at 72°C. The 2nd round PCRs were run with the inner primer sets (Table 1) and 1 µl of the 1st round products and used the same cycling conditions as the 1st round. The resulting products were evaluated on a 1% agarose gel and wells with the expected band sizes (∼600 base pairs (bp) for the heavy chain and ∼550 bp for the light chains; Supplemental Fig. 1) were excised and purified using QIAquick Gel Extraction kits per the manufacturer’s instructions. The purified samples were Sanger sequenced by Azenta Life Sciences using the 3’ inner Ig constant domain primers. The sequences were analyzed using Geneious Prime and the online IMGT/HighV-QUEST tool. The productive heavy and light chain pairs were prepared for antibody production.

### 2.5 Cloning Ig variable fragments into expression vectors

For each Ig molecule, we used the nested PCR product sequences to select gene-specific 5’ and 3’ cloning primers. These primers were adapted from Sundling et al. (Sundling et al., 2012b) and Magnani et al. (Magnani et al., 2017) and contain restriction sites (Heavy chain: 5’ EcoRI, 3’ NheI; Kappa chain: 5’ EcoRI, 3’ BstAPI; and Lambda chain: 5’ EcoRI, 3’ XhoI) that are compatible with cloning into the rhesus IgG1 pFUSE expression vectors (pFUSEss-CHIg-rhG1 and pFUSEss-CLIg-rhK from Invivogen; rhIgLC1*01 courtesy of D. Magnani, (Magnani et al., 2017)). The cloning PCR mix consisted of 1 µl of the nested PCR product, 12.5 µl of 2X Phusion HF master mix (ThermoFisher Scientific), 1.25 µl of 500 nM forward and reverse cloning primers, and 9 µl of nuclease-free water. The PCR program was as follows: 98°C for 30 seconds, 25 cycles of 98°C for 10 seconds, 58°C for 20 seconds for light chains or 62°C for 20 seconds for heavy chains, 72°C for 15 seconds, a final elongation step of 72°C for 5 minutes, followed by cooling to 10°C. A 5 µl aliquot of each PCR reaction was run on a 1% agarose gel to confirm the presence of an amplification product of the correct size. The remaining PCR products were purified using the QIAquick PCR purification kit.

Next, we digested the purified PCR products using 3 µl of 10X Cutsmart buffer, 23 µl of PCR product, 0.5 µl of EcoRI, 0.5 µl NheI (heavy chain) or 0.5 µl of BstAPI (kappa chain) or XhoI (lambda chain), and 3 µl of nuclease-free water. All restriction enzymes were from New England BioLabs. Following digestion, the PCR products (∼450 bp) were purified using the QIAquick PCR purification kit and ligated into the appropriate linearized expression vectors using T4 DNA ligase (New England BioLabs) and incubated overnight at 16°C.

Finally, the ligated vectors were transformed into NEB 5-alpha chemically competent E. coli (New England BioLabs) according to the manufacturer’s protocol and incubated overnight at 37°C on LB agar plates containing zeocin (heavy chains) or blasticidin (light chains). Select colonies were then grown in 3 ml of LB with appropriate antibiotics and the plasmids were purified with PureYield Plasmid Miniprep System (Promega), performing restriction enzyme digests to verify that the plasmids had inserts.

### 2.6 Expression of recombinant mAbs

Recombinant mAbs were transiently produced in FreeStyle 293F cells (ThermoFisher Scientific) following the protocol detailed by Vink *et al* (Vink, Oudshoorn-Dickmann, Roza, Reitsma, & de Jong, 2014). First, the cells were suspended in FreeStyle 293 Expression Medium (ThermoFisher Scientific) at a concentration of 1.1×10^6 cells/ml, and 2 ml of the suspension were added to individual wells of a 24-well deep well plate. The cells were transfected with a 2 µg mixture of plasmids encoding the heavy chain (690 ng), light chain (690 ng), human p21 (Invivogen; 100 ng), human p27 (Invivogen; 500 ng), and SV40 large T antigen (20 ng) suspended in Opti-MEM I Reduced-Serum Medium (ThermoFisher Scientific), mixed with 293fectin (ThermoFisher Scientific), and added to the cells per the manufacturer’s instructions. The p21-, p27-, and SV40 large T antigen-encoding plasmids were included to enhance antibody production, as previously described (Vink et al., 2014). After transfecting the cells, the plate was clamped into a Microflask system (Applikon Biotechnology) affixed to an orbital shaker (Thermo Scientific) in a 37°C, 5% CO2 incubator. The supernatant was harvested after 7 days.

### 2.7 SIV Env ELISA

We tested the specificity of the mAbs in an enzyme-linked immunosorbent assay (ELISA). A 2HB 96-well plate (Immulon) was coated overnight at 4°C with 100 µl of a 5 µg/ml solution of SIVmac239 Env gp130. The following day, the antigen was removed, and the plate was blocked with blocking buffer (PBS + 2% BSA) for 1 hour at room temperature. The FreeStyle 293F supernatants were diluted 1:10 in blocking buffer and then serially diluted 1:4 until a dilution factor of 10,240 was achieved. After removing the blocking buffer, the plate was washed six times with wash buffer (PBS + 0.5% tween-20) and then 100 µl of the serially diluted supernatant aliquots were added to the plate, followed by a 2-hour incubation at 37°C. A similarly prepared anti-MHC antibody (anti-Mamu-A1*001) was used as a negative control (Holman et al., 2017). Subsequently, the plate was washed six times with wash buffer, and 100 µl of a 1:1000 dilution of horseradish peroxidase-conjugated anti-macaque IgG1/3 antibody (1B3; NIH Nonhuman Primate Reagent Resource) was added to each well. After a 1-hour incubation at room temperature, the plate was washed six times with wash buffer and 100 µl of TMB substrate solution (ThermoFisher Scientific) was added to the wells. After being incubated for ∼30 minutes at room temperature, the reaction was terminated with 100 µl of 1N HCL. The optical density of each well was determined with a GloMax-Multi Detection System (Promega) reading at 450 nm.

## Results

### 3.1 Single-cell sorting and rescue of antibody sequences

To demonstrate the cloning process, we isolated SIV Env-specific antibodies from a rhesus macaque chronically infected with SIVmac239 (Mudd et al., 2012). Using cryopreserved PBMCs, we isolated SIV Env-binding memory B cells using biotinylated SIV Env gp130 as a bait antigen and labeled cell-bound antigens with PE-conjugated streptavidin (Fig. 1), as previously described (Mohanram et al., 2014). Thus, we sorted single class-switched SIV-Env-binding memory B cells (live, CD3-, CD14-, CD16-, CD20+, CD27+, IgM-, SIV Env gp130+) into individual wells of a 96-well plate containing lysis buffer and poly(dT) RT oligos (Table 1). The poly(dT) RT oligo has a 30 deoxythymidine (dT) tract suitable for 3’ RACE, annealing to mRNA poly-A tails to initiate cDNA synthesis during reverse transcription, and a unique 22 bp sequence at the 5’ end, creating the 3U synthetic primer binding site.

We attempted to rescue the heavy and light chain variable regions from 14 sorted cells. The first step was generating full-length cDNA from mRNA in RT reactions. We want to highlight several components of the RT mix. First, we used Superscript IV, a thermostable MMLV-based RT with high processivity and transferase activity (Zucha, Androvic, Kubista, & Valihrach, 2020), allowing for the RT reactions to be performed at a higher temperature (50°C) than most cDNA synthesis reactions (42°C), helping to destabilize mRNA secondary structure and produce full-length cDNA (Arezi & Hogrefe, 2009; Baranauskas et al., 2012). Second, we included betaine, an N-trimethylated amino acid that interferes with G–C nucleotide pairs, to further destabilize the secondary structure of mRNAs (Picelli et al., 2013; Pinto & Lindblad, 2010; Rees, Yager, Korte, & von Hippel, 1993; Spiess & Ivell, 2002). Third, the template switching oligo (TSOUoUi) contains three 3’-terminal riboguanosines (rG), which improves its binding to the non-templated cytosines added by Superscript IV to the 3’ end of nascent cDNAs (Zhu et al., 2001). Notably, TSOUoUi contains two synthetic primer-binding sites, 5UO and 5Ui, that permit nested PCRs with the corresponding “universal” primers. The 5UO primer is identical to the ISPCR primer described in Picelli et al. (Picelli et al., 2014) and the 5Ui primer is a scrambled version of the ISPCR. In sum, the RT reactions produce full-length cDNAs with synthetic primer binding sites appended to both ends, enabling PCR amplification with universal primers.

Recovering heavy and light chain pairs from memory B cells can be challenging due to the low abundance of BCR transcripts. To address this issue, we included a preamplification step using the 5UO and 3U primers to increase the amount of cDNA available for subsequent steps. We then performed nested PCRs to amplify the Ig heavy and light chain V genes using the 5UO and 5Ui primers along with gene-specific primers targeting the IgGH, Igκ, or Igλ constant domains. Additionally, we used touchdown cycling conditions in the nested PCRs to reduce nonspecific amplifications. Importantly, the 5UO and 5Ui primers enabled the amplification of the IgV genes without the need for diverse 5’ MTPX primer sets.

After the second round of amplification, the PCRs yielded ∼550-600 bp amplicons (Supplemental Fig. 1) that were gel purified and Sanger sequenced. We used the sequencing results to determine the specific V gene encoded by each Ig molecule and to select gene-specific cloning primers. These primers included restriction sites to insert the PCR products into antibody expression plasmids. Using this strategy, we recovered 10 heavy and light chain pairs from 14 cells.

### 3.2 Antibody expression and binding

To produce the recombinant antibodies, we first transformed E. coli with the plasmids containing the cloned IgV genes and screened for bacterial clones containing inserts of expected sizes, ∼500 bp for heavy chains and ∼450 bp for light chains. The recombinant mAbs were produced by co-transfecting HEK293F cells with paired heavy and light chain expression plasmids. After incubating the transfected cells for 7 days, we tested the specificity of the cloned antibodies using serial dilutions (1:10 to 1:10,240) of the supernatants in SIV Env gp130 ELISAs (Fig. 3). Supernatant from HEK293F cells transfected with plasmids encoding a mAb recognizing an irrelevant epitope was used as a negative control. Eight of the ten cloned mAbs bound to SIV Env gp130, demonstrating this method’s ability to rescue mAbs from individual RM antigen-specific memory B cells (10 mAbs from 14 sorted cells) that recognize their cognate antigen (8 of 10 mAbs).

**Figure 3.**
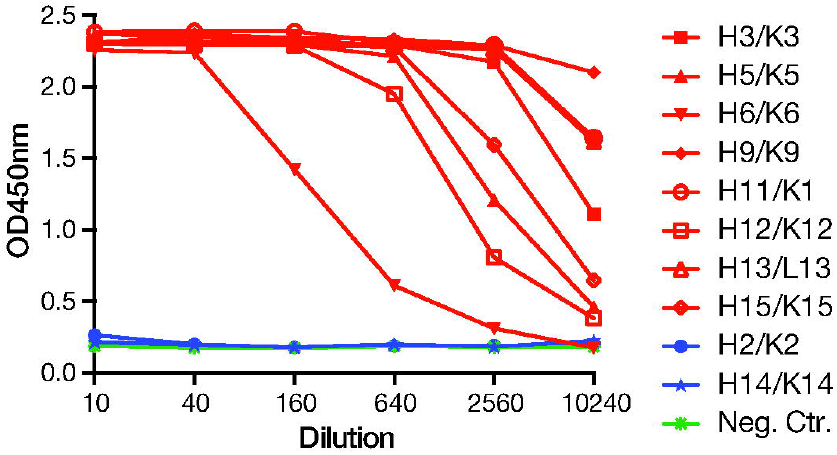
Binding of mAbs to SIV Env gp130. Anti-SIV Env gp130 ELISA titration curves for supernatants from HEK293F cells transfected with plasmids encoding 10 recombinant mAbs. mAbs showing optical density (OD) 450 above 1 at a 1:160 dilution are highlighted in red. A mAb binding an irrelevant epitope was used as a negative control (green).

### 3.3 V(D)J gene usage of cloned mAbs

Characterizing Ig gene profiles and determining the extent of somatic hypermutation is an integral component of pre-clinical studies in RM. Thus, we used the IMGT RM germline database and V-QUEST alignment software to analyze the V(D)J gene usage and somatic hypermutation levels in the isolated mAbs (Table 2). All of the antibodies used VH genes from either the VH3 or VH4 family, consistent with previous studies finding these are the predominant IgH gene families used by rhesus macaques (Dai et al., 2015; Silveira et al., 2015; Sundling et al., 2012a; Sundling et al., 2012b; Sundling et al., 2014; Chernyshev et al., 2021; Vázquez Bernat et al., 2021). Nine of the mAbs used κ light chains. Notably, four of the mAbs (H6/K6, H12/K12, H14/K14, and H15/K15) used the same VH allele (*IGHV3-28*02*), but they were not clonally related as they used different *IGVD* and *IGVJ* genes. However, two of them (H6/K6 and H12/K12) were paired with a κ light chain using the same *IGKV* allele (*IGKV-104*01*) but with different IGKJ genes. Of the mAbs using *IGHV3-28*02*, all bound to SIV Env except H14/K14. Additionally, the heavy chains had an average of 17 amino acid changes with a range of 4-27 mutations, while the light chains had an average of 10 amino acid changes with a range of 3-17 amino acid mutations. These results show that the cloning process presented here can be used to characterize Ig genes in individual memory B cells.

**Table 2.** Genetic characterization of the isolated mAbs.

| mAb | Heavy chain |  |  | CDR3<br>length (aa) | Number of<br>Mutations<br>(aa) | Light chain |  | CDR3<br>length (aa) | Number of<br>Mutations<br>(aa) |
| --- | --- | --- | --- | --- | --- | --- | --- | --- | --- |
|  | IGHV | IGVD | IGVJ |  |  | IGV | IGJ |  |  |
| H2/K2 | IGHV3-136*01 | IGHD3-16*01 | IGHJ1*01 | 12 | 17 | IGKV2-86*01 | IGKJ4*01 | 9 | 9 |
| H3/K3 | IGHV3-8*01 | IGHD4-35*01 | IGHJ4*01 | 17 | 9 | IGKV1-33*01 | IGKJ2*01 | 9 | 8 |
| H5/K5 | IGHV3-54*01 | IGHD6-31*01 | IGHJ6*01 | 14 | 17 | IGKV2-78*01 | IGKJ3*01 | 9 | 17 |
| H6/K6 | IGHV3-28*02 | IGHD2-39*02 | IGHJ5-2*02 | 19 | 27 | IGKV2-104*01 | IGKJ3*01 | 10 | 8 |
| H9/K9 | IGHV3-59*01 | IGHD2-39*02 | IGHJ4*01 | 15 | 25 | IGKV2S8*01 | IGKJ4*01 | 8 | 8 |
| H11/K11 | IGHV4-127*01 | IGHD3-34*01 | IGHJ4*01 | 11 | 23 | IGKV2-65*01 | IGKJ2*01 | 10 | 8 |
| H12/K12 | IGHV3-28*02 | IGHD6-25*01 | IGHJ6*01 | 15 | 4 | IGKV2-104*01 | IGKJ4*01 | 9 | 3 |
| H13/L13 | IGHV4-160*01 | IGHD4-29*01 | IGHJ5-1*02 | 18 | 19 | IGLV3-25*02 | IGLJ6*01 | 11 | 15 |
| H14/K14 | IGHV3-28*02 | IGHD3-16*01 | IGHJ6*01 | 17 | 5 | IGKV2-99*01 | IGKJ2*01 | 9 | 17 |
| H15/K15 | IGHV3-28*02 | IGHD1-1-1*01 | IGHJ5-2*01 | 13 | 21 | IGKV1-38*02 | IGKJ3*01 | 10 | 11 |

### 3.4 Comparison of Ig leader sequences to established nested PCR primer sets

The nested PCR primer sets developed by Sundling et al. (Sundling et al., 2012b) are commonly used to isolate mAbs from single RM B cells (Silveira et al., 2015; Sundling et al., 2014; Magnani et al., 2017; Li, Hessell, Kong, Haigwood, & Gorny, 2021; Chen et al., 2021; Moin et al., 2022; van Schooten et al., 2021; Yacoob et al., 2018; Feng et al., 2023). These primer sets, designated L1 (outer) and SE (inner), consist of 9-14 primers that anneal to two distinct regions of the V gene leader sequences (Sundling et al., 2012b). However, these primer sets may not capture the full spectrum of RM IgV leader sequence diversity, leading to decreased cloning efficiency. Since the nested PCRs described here yielded unbiased amplification of the IgV leader sequences, we assessed the conservation of the Sundling L1 and SE primers in the leader sequences of the 10 heavy and light chain pairs.

Of the isolated antibodies, only one (H2/K2) had leader sequences that perfectly matched at least one of the L1 and SE primers for both the heavy and light chains (Supplemental Fig. 2). Furthermore, only four individual Ig chains had leader sequences that perfectly matched a primer in the outer L1 and inner SE set. Notably, seven of the IgV leader sequences had at least one nucleotide mismatch with all of the L1 and SE primers. Although some of the L1 and SE primers would have annealed despite these nucleotide mismatches, these findings highlight the challenges of amplifying RM IgV genes using nested PCR primers that bind to Ig leader sequences and likely contribute to the relatively low efficiency of isolating paired Ig heavy and light chains (Sundling et al., 2014; Silveira et al., 2015).

## 4.Discussion

Here we present a method for efficiently isolating mAbs from RMs that addresses several challenges in rescuing antibodies from individual memory B cells. The key feature of this process is combining single-cell RNA-seq methods with established mAb cloning procedures. To isolate IgV genes, we leverage 3’RACE and the intrinsic terminal transferase activity of an MMLV-based RT to append synthetic primer binding sites to the 5’ and 3’ ends of cDNA during synthesis. As a result, this approach (1) allows for the amplification of low abundance antibody templates and (2) eliminates the need for degenerate MTPX primers to improve the recovery of heavy and light chain pairs from antigen-specific RM memory B cells.

A core element of this protocol is using 5’ RACE to generate the cDNA libraries from individual sorted cells. 5’ RACE is a well-established method for characterizing the 5’ end of mRNA transcripts and forms the basis for the SMART cDNA synthesis technology (Zhu et al., 2001). However, traditional 5’ RACE techniques are not suitable for synthesizing cDNA from single cells due to the small quantities of mRNA available. Recent advancements in single-cell RNA sequencing technologies and the development of MMLV-based RTs with improved thermostability and processivity have improved cDNA library synthesis from single cells (Picelli et al., 2013; Picelli et al., 2014; Rogers & Potter, 2017; Zucha et al., 2020). To adapt this technology to cloning RM antibodies, we applied principles from the SMART-Seq2 protocol (Picelli et al., 2013; Picelli et al., 2014) to prepare cDNA libraries from individual RM memory B cells. Although, we performed the cDNA synthesis reactions at 50°C instead of 42°C as, in our experience, it was necessary to generate full-length products. Importantly, adapting these RNA-seq techniques allowed us to add universal primer binding sites to the 5’ ends of cDNAs, reducing the number of 5’ primers used in each nested PCR reaction to a single unique forward primer.

Previous reports have described 5’ RACE-based strategies for cloning mAbs. The first published study used Superscript III, a thermostable MMLV-based RT, to synthesize cDNAs at 55°C (Ozawa, Kishi, & Muraguchi, 2006). Yet, this RT has poor transferase activity, so a subsequent tailing step with terminal deoxynucleotidyl-transferase (TdT) was necessary to add non-templated nucleotides to the cDNA. However, TdT adds nucleotides to both incomplete and full-length cDNAs, leading to the amplification of non-specific products. In contrast, we use Superscript IV that, in a single step, synthesizes full-length cDNA and adds non-templated nucleotides to the 3’ end of nascent cDNAs. Recently, several groups have employed single-cell SMART-seq-based approaches to characterize IgV genes from individual B cells using next-generation sequencing and specialized analysis pipelines to (Neu et al., 2019; Upadhyay et al., 2018; Zost et al., 2020), including the BALDR pipeline to reconstruct RM Ig sequences (Upadhyay et al., 2018). Nevertheless, these approaches require bioinformatic expertise to recover the Ig sequences and use existing primer sets to amplify IgV genes from cDNA. Therefore, unique aspects of our cloning procedures are using SMART 5’ and 3’ RACE reactions to append synthetic primer binding sites to the 5’ end of cDNAs during synthesis and using universal primers in nested PCRs to amplify IgV genes.

Although we focused on cloning antibodies from RM memory B cells, we anticipate this method will be adaptable to other antigen receptors. For example, this technique may capture RM T cell receptor (TCR) sequences from individual T cells by substituting the Ig constant region primers for primers complementary to TCR alpha and beta chains. Likewise, this method may be useful for isolating antibodies from species with poorly characterized IgV genes (ex. ferrets and Syrian golden hamsters), as only prior knowledge of Ig constant region sequences is required to design gene-specific primers for the nested PCRs while cloning primers can be generated from the V gene sequences.

In this study, we sorted memory B cells using SIV Env probes, but antigenic probes are not always available to isolate antigen-specific B cells. However, we expect our approach will also be useful for cloning antibodies from plasmablasts, given these cells typically have abundant antibody transcripts (Magnani et al., 2017; Silveira et al., 2015). Moreover, some antigens elicit numerous binding antibodies, but only a few with desired effector functions (i.e., neutralizing activity). Therefore, we believe that our method can be paired with short-term culturing or immortalization techniques to screen large numbers of antibody-secreting cells for desired properties (Blasi et al., 2018; Huang et al., 2013; Zhao et al., 2020) before isolating mAbs from the selected cells.

## 5. Conclusion

In summary, we demonstrate a SMART-based method for efficiently isolating mAbs from RMs. This process involves incorporating synthetic primer binding sites into cDNAs during synthesis, allowing for unbiased amplification of IgV genes using universal 5’ primers in nested PCR reactions. This process may be adaptable to isolating RM TCRs and mAbs from other animal species with minimal modifications. We anticipate this approach will be a valuable tool for characterizing antibody functions and IgV gene usage in vaccine and infectious disease studies.

## Abbreviations

RM: rhesus macaque
mAbs: monoclonal antibodies
SMART: switching mechanism at the 5’ ends of the RNA transcript
IgV: immunoglobulin variable
MTPX: multiplex
FACS: fluorescent-activated cell sorting
RACE: rapid amplification of cDNA ends
RT: reverse transcriptase
dT: deoxythymidine
TSO: template-switching oligo
SIV: simian immunodeficiency virus
PBMC: peripheral blood mononuclear cells
TdT: terminal deoxynucleotidyl-transferase
BCR: B-cell receptor
TCR: T-cell receptor

## Acknowledgments

We thank Dr. Diogo Magnani for his helpful discussions. This study was supported by National Institutes of Health (NIH) grant R37AI095098 and contract 75N93021C00006.

Additional support was provided by the Office of Research Infrastructure Programs/OD via grant P51OD011106 awarded to the WNPRC at the University of Wisconsin-Madison. The funders had no role in study design, collection, analysis, interpretation of the data, or manuscript preparation. The content is solely the responsibility of the authors and does not necessarily represent the official views of the NIH. The authors have no competing interest to declare.

The following reagent was obtained through the NIH HIV Reagent Program, Division of AIDS, NIAID, NIH: Simian Immunodeficiency Virus (SIV) SIVmac239 gp130-His Recombinant Protein, ARP-12797, contributed by Dr. Klaus Uberla. The Anti-rhesus IgG1/3 [1B3] antibody used in this study was provided by the NIH Nonhuman Primate Reagent Resource (NIAID U24 AI126683).

