## Supplemental Figure 1 for "A SMART method for efficiently isolating monoclonal antibodies from individual rhesus macaque memory B cells"

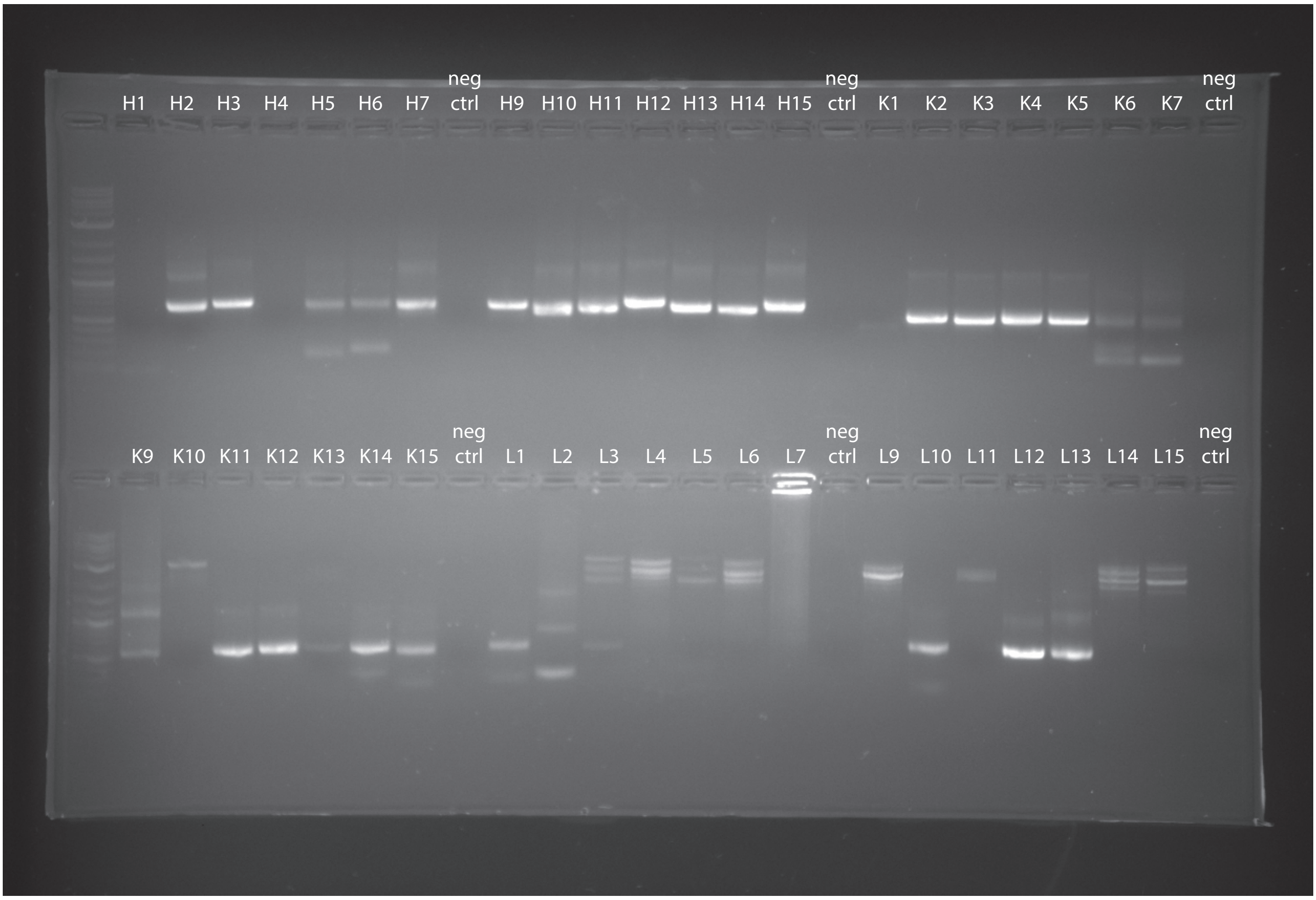


**Supplemental Figure 1.** PCR products after the second round amplifications run on a 1% agarose gel. The heavy (H1-H15) chain yields ~600pb bands and the kappa (K1-K15) and lambda (L1-L15) chains yield ~550bp bands.
