## Supplemental Figure 2 for "A SMART method for efficiently isolating monoclonal antibodies from individual rhesus macaque memory B cells"

```
H2       ATGGAGTTGGGGCTGAGCTGGGTTTTCCTTGTTGCTATTTTAAAAGGTGTCCAGTGTG
VH3A.L1  ATGGAGTTKGGGCTGAGCTG
VH3E.SE                                   GCTATTTTAAAAGGTGTCCAGTGTG

H3       ATGGAATTTAGGCTGAGCTGGGTTTTCCTTGTTGGTGTTTTAAAAGGTGTCCAGTGTG
VH3A.L1  ATGGAGTTKGGGCTGAGCTG
VH3E.SE                                   GCTATTTTAAAAGGTGTCCAGTGTG

H5       ATGAAGTTTGGGCTGGGCTGGGTTTTCCTTGTTGCCATTTTGAGAGGTGTCCAGTGTG
VH3A.L1  ATGGAGTTKGGGCTGAGCTG
VH3D.SE                                   GCTATTTTAAGAGGTGTCCAGTGTG

H6       ATGGAGTTTGTGCTGAGCTGGGTTTTCCTGGTTGCTCTTTTGAAAGGGGTCCAGTGTG
VH3B.L1  ATGGAGTTTGKRCTGAGCTGG
VH3C.SE                                   GCTCTTTTGAAAGGTGTCCAGTGTG

H9       ATGGAGTTTGTGCTGAGCTGGGTTTTCCTTGTTGCTCTTTTGAAAGGTGTCCAGTGTG
VH3B.L1  ATGGAGTTTGKRCTGAGCTGG
VH3C.SE                                   GCTCTTTTGAAAGGTGTCCAGTGTG

H11      ATGAAGCACCTGTGGATCTTCCTTNTCCTGGTGGCAGCTCCCAGATGGGTCCTGTCC
VH.L1    ATGAAGCACCTGTGGTTC
VH.SE                                       AGCTCCCAGATGGGTCYTGTCC

H12      ATGGAGTTTGGCCTAAGCTGGGTTTTCCTTGTTGCTATTTTAAAAGGTGTCCAGGGCG
VH3A.L1  ATGGAGTTKGGGCTGAGCTG
VH3E.SE                                   GCTATTTTAAAAGGTGTCCAGTGTG

H13      ATGAAGCACCTGTGGTTCTTCCTCCTCCTGGTGGTAGCTCCCAGATGGGTCCTGTCC
VH4.L1   ATGAAGCACCTGTGGTTC
VH4.SE                                      AGCTCCCAGATGGGTCYTGTCC

H14      ATGGAGTTTGGCCTAAGCTGGGTTTTCCTTGTTGCTATTTTAAAAGGTGTCCAGGGTG
VH3A.L1  ATGGAGTTKGGGCTGAGCTG
VH3E.SE                                   GCTATTTTAAAAGGTGTCCAGTGTG

H15      ATGGAGTTTGGCCTAAGCTGGGTTTTCCTTGTTGCTATTTTAAAAGGTGTCCAGGGTG
VH3A.L1  ATGGAGTTKGGGCTGAGCTG
VH3E.SE                                   GCTATTTTAAAAGGTGTCCAGTGTG

K2       ATGAAGCTCCCTGCTCAGCTCCTGGGGCTGCTAATGCTCTGGCTCCCTGGATCCAGTGGG
VK2.L1   ATGARGYTCCCTGCTCAG
VK1B.SE                                          GGTCCCTGGRTCCAGTGGG

K3       ATGGACATGAGGGTCCTCACTCAGCTCCTGGGGCTCCTGCTGCTCTGGCTCCCAGGTGCCAGATGTGA
VK1A.L1  ATGGACATGAGGGTCCCCGC
VK1/2.SE                                                CTCCCAGGTGCCAGATGTGA

K5       ATGAGGCTCCCTGCTCAACTCCTGGGGCTACTGGTGCTCTGGCTCCCTGGGACCACTGGA
VK2.L1   ATGARGYTCCCTGCTCAG
VK3A.SE                                        TGGCTCCCAGGTACCACYGGA

K6       ATGAGGCTCCCTGCTCAGCTCCTGGGACTGCTAATGCTCTGGCTCCCTGGATCCAGTGGG
VK2.L1   ATGARGYTCCCTGCTCAG
VK1B.SE                                          GGTCCCTGGRTCCAGTGGG
```

```
K9       ATGAGGCTCCCTGCTCAGCTCCTGGGGCTGCTATTGCTCTGCGTCCCCGGATCCAGTGGG
VK2.L1   ATGARGYTCCCTGCTCAG
VK1B.SE                                           GGTCCCTGGRTCCAGTGGG

K11      ATGAGGCTCCCTGCTCAGCTCCTGGGGCTGCTGTTGCTCTGCGTCCCCGGGTCCAGTGGG
VK2.L1   ATGARGYTCCCTGCTCAG
VK1B.SE                                           GGTCCCTGGRTCCAGTGGG

K12      ATGAGGCTCCCTGCTCAGCTCCTGGGACTGCTAATGCTCTGGCTCCCTGGATCCAGTGGG
VK2.L1   ATGARGYTCCCTGCTCAG
VK1B.SE                                           GGTCCCTGGRTCCAGTGGG

L13      Unavailable


K14      ATGAGGCTCCCTGCTCAGCTCCTGGGGCTGCTGGTGCTCTGGCTCCCTGGGACCAGTGGG
VK2.L1   ATGARGYTCCCTGCTCAG
VK1B.SE                                           GGTCCCTGGRTCCAGTGGG

K15      ATGGACATCAGGGTCCTCGCTCAGCTTCTCGGGCTCCTGCTGCTCTGGCTCCCAGGTGCCACATGTGA
VK1A.L1  ATGGACATGAGGGTCCCCGC
VK1/2.SE                                                 CTCCCAGGTGCCAGATGTGA
```

**Supplemental Figure 2. Comparison of Ig leader sequences to commonly used nested PCR primers for amplifying RM IgV genes.** We compared the leader sequences from the heavy and light chain pairs for the 10 isolated mAbs to the most closely related outer (L1) and inner (SE) primers described in Sundling *et al*.[17] Nucleotide mismatches between the primer and leader sequences are highlighted in red and underlined.
